# Dynamic connectivity analyses of intracranial EEG during recognition memory reveal various large-scale functional networks

**DOI:** 10.1101/2020.10.30.360529

**Authors:** Jakub Kopal, Jaroslav Hlinka, Elodie Despouy, Luc Valton, Marie Denuelle, Jean-Christophe Sol, Jonathan Curot, Emmanuel J. Barbeau

## Abstract

Recognition memory is the ability to recognize previously encountered events, objects, or people. It is characterized by its robustness and rapidness. Even this relatively simple ability requires the coordinated activity of a surprisingly large number of brain regions. These spatially distributed, but functionally linked regions are interconnected into large-scale networks. Understanding memory requires an examination of the involvement of these networks and the interactions between different regions while memory processes unfold. However, little is known about the dynamical organization of large-scale networks during the early phases of recognition memory. We recorded intracranial EEG, which affords high temporal and spatial resolution, while epileptic subjects performed a visual recognition memory task. We analyzed dynamic functional and effective connectivity as well as network properties. Various networks were identified, each with its specific characteristics regarding information flow (feedforward or feedback), dynamics, topology, and stability. The first network mainly involved the right visual ventral stream and bilateral frontal regions. It was characterized by early predominant feedforward activity, modular topology, and high stability. It was followed by the involvement of a second network, mainly in the left hemisphere, but notably also involving the right hippocampus, characterized by later feedback activity, integrated topology, and lower stability. The transition between networks was associated with a change in network topology. Overall, these results confirm that several large-scale brain networks, each with specific properties and temporal manifestation, are involved during recognition memory. Ultimately, understanding how the brain dynamically faces rapid changes in cognitive demand is vital to our comprehension of the neural basis of cognition.

- Various dynamic large-scale networks support recognition memory.
- The first is mostly feedforward and involves the right hemisphere and the bilateral frontal lobes.
- The second is mostly feedback and includes left MTL regions and the right hippocampus.
- Changes in network topology accompany the switch between the networks.

## 1. Introduction

Visual recognition memory has been studied since the 1960s (Milner, 1972) with a tremendous number of findings that have helped to reveal how massively accurate (Brady et al., 2008), fast (Besson et al., 2012), and long-lasting (Larzabal et al., 2018) it can be. The remarkable efficiency and robustness of this system imply that it has a strong ecological value. These studies have also pinpointed the medial temporal lobes as critical for this type of memory (Brown & Aggleton, 2001; Eichenbaum et al., 2007). In a broader sense, it has consistently been shown that visual recognition memory relies on the “what” system, i.e., the visual ventral stream, which involves many temporo-basal brain regions such as the lingual, fusiform, and parahippocampal gyri. The participation of the ventral stream is asymmetric in the sense that visual recognition memory relies more on the right than on the left hemisphere (Milner & Taylor, 1972; Patterson & Bradshaw, 1975; Elger et al., 1997; Barbeau et al., 2008). In addition, visual recognition memory also involves parietal and frontal lobe regions, probably for processes concerned with confidence and decision-making (for a review and a model, see Bastin et al. 2019).

Even a relatively simple task, such as deciding whether an object has already been seen or not, thus requires the involvement of a surprisingly large number of brain regions. The temporal dynamics of recognition are now better understood as the first behavioral responses occur in approximately 360 ms (Besson et al., 2012), the first neural differences between targets and distractors are identified at approximately 200 ms (Barbeau et al., 2008; Caharel et al., 2014; Barragan-Jason et al., 2015) and many different brain regions are involved up to 600 ms or more. Even though the activity of participating brain regions appears to be partly sequential, it is mostly overlapping (Despouy et al., 2020), and what specific interactions take place between regions is unknown.

Brain regions do not operate in isolation but are interconnected in large-scale networks (Varela et al., 2001; Betzel et al., 2019; Bressler & Menon, 2010; De Luca et al., 2006). The basis of every network is connectivity, defined as either anatomical links (structural connectivity), statistical dependencies (functional connectivity), or causal interactions (effective connectivity) (Sporns, 2007). Substantial evidence supports the hypothesis that the architecture of brain networks is non-random and is optimized to support cognitive abilities. Interesting properties underpin this efficient architecture, such as high modularity (see Bassett & Sporns 2017). The modular architecture is characterized by small subsystems (communities), composed of different brain regions with a vast number of local connections and few distant connections. This hierarchically modular structure supports effective communication (Lynn et al., 2020) as well as functional segregation and specialization (Sporns, 2013).

Given the continually evolving environment, and depending on the system’s demands, there are continuously changing patterns of interactions between brain regions (Sporns, 2013; Bassett et al., 2011). Therefore, both the topology of the networks and the interactions between them are highly dynamic (Allen et al., 2014; Shine et al., 2016; Zalesky et al., 2014; Hutchison et al., 2013). As a result, it has been suggested that dynamic network reconfiguration is a fundamental neurophysiological process (Braun et al., 2015; Kitzbichler et al., 2011; Vatansever et al., 2015). Emerging findings suggest that networks are non-stationary (see Hutchison et al. 2013), although robust characterization of this non-stationarity remains a methodological challenge (Hlinka & Hadrava, 2015; Kaplan et al., 2005). It is generally assumed that their reconfigurations are driven by higher-order cognitive control systems, involving mainly the frontal cortex (Fedorenko & Thompson-Schill, 2014; Braun et al., 2015). Moreover, dynamic reconfiguration is directly linked to cognitive performance during memory (Cohen & D’Esposito, 2016; Stevens et al., 2012; Meunier et al., 2014; Stanley et al., 2014).

Although it is clear that recognition memory requires the participation of different networks, little is known about their dynamical organization. This is due to the fact that current studies of large-scale network dynamics are based either on fMRI or EEG. fMRI-based temporal networks are usually analyzed using multiple, possibly overlapping, very long temporal segments, typically 30-60 seconds long (Allen et al., 2014; Hutchison et al., 2013). In contrast, surface EEG studies suffer from low spatial resolution and might not capture the contributions of medial temporal brain structures.

Because visual recognition memory is so fast, the modifications of large-scale functional networks that support such ability need to be examined on a millisecond-by-millisecond timescale and with high spatial resolution. Therefore, we analyzed intracranial EEG, an approach that meets these needs. We calculated functional and effective connectivity, as well as underlying graph properties. Considering that the contribution to visual recognition memory of each hemisphere differs significantly, we assumed that it would be reflected in the connectivity patterns. We ran the first set of analyses based on this hypothesis. We then examined whole-brain network topology and investigated fluctuations in network properties, i.e., changes in integration and segregation as memory processes unfold (Cohen & D’Esposito, 2016; Vatansever et al., 2015). Ultimately, understanding how brains dynamically adapt to perform very fast tasks is vital to our understanding of the neural basis of memory.

## 2. Material and Methods

### 2.1. Patients

Intracranial EEG (iEEG) was recorded for eighteen patients with drug-refractory epilepsy (8 women, age: 37.61 ± 11.37 years old). They were admitted to the Epilepsy Monitoring Unit at Toulouse University Hospital for the identification and possible subsequent resection of the epileptogenic zone. Eight – thirteen depth electrodes were stereotaxically implanted in each patient. The depth electrodes were 0.8 mm in diameter and contained 8 to 18 platinum/iridium contacts, each 2 mm long (Microdeep depth electrode, DIXI medical, France). Each implantation was individually tailored to the seizure onset zone and the placement of each depth electrode was based exclusively on clinical criteria independently of this study.

The preoperative MRI and postoperative CT images were fused and normalized to the Montreal Neurological Institute (MNI) brain atlas for precise contact localization (see Despouy et al. 2020).

Intracranial EEG activity was recorded using two synchronized 64-channel acquisition systems (SystemPlus Evolution, SD LTM 64 EXPRESS, Micromed, France) with a sampling frequency of 256 Hz for two patients and either 1,024 or 2,048 Hz for the others (high pass-filter: 0.15 Hz). None of the patients had a seizure within 6 hours before the recordings.

This study was approved by the local University Hospital Ethics Committee (CER No. 47–0913). Informed consent forms were signed for the implantation and the use of iEEG data for research purposes.

### 2.2. Visual recognition memory test

Each subject performed a visual recognition memory task, namely the Speed and Accuracy Boosting procedure (SAB) while the intracerebral EEG was being recorded (Besson et al., 2012). Each block began with an encoding phase during which 30 trial-specific stimuli (targets) were presented individually for at least 3 s (self-paced) in the center of a gray screen. The stimuli were taken from an extensive database of high-quality cropped photos of everyday objects. Participants were explicitly instructed to remember all stimuli. A distracting phase followed during which the subjects watched a colored cartoon video with sound on for 3 minutes. Finally, the subjects underwent the recognition memory phase during which the 30 targets and 30 distractors were shown. Subjects were required to respond to the targets only, by raising their finger as quickly as possible from an infrared pad. A 600ms response time limit with audio feedback forced subjects to answer as quickly and accurately as possible. Responses were based on a go/no-go design. If a go response was given before the response time limit, positive audio feedback was played if the stimulus was a target (Hit). Negative feedback was played if it was a distractor (False alarm). If a no-go response was given, positive audio feedback was played if the stimulus was a distractor (Correct rejection). Negative feedback was played if the stimulus was a target (Miss). We analyzed only the Hits and Correct rejections (CR) in this study. The SAB test is demanding and requires one or two training sessions, which were not included in subsequent analyses. Patients participated in 7–10 SAB blocks depending on their willingness.

We evaluated each subject’s performance using two discrimination indices, i.e., d-prime and minimal reaction time (minRT). The minRT is defined as the minimal processing time required to recognize targets, and it was computed by determining the latency at which correct go responses (Hits) started to significantly outnumber incorrect go responses (false alarms) (Besson et al., 2012). As in previous studies (Despouy et al., 2020), we used 20ms time bins and a Fisher’s exact test (*p* < 0.05), followed by at least two significant consecutive bins to compute the minRT.

### 2.3. Recordings

We used a bipolar montage between adjacent contacts to remove artifact contaminations, identify local activations, and provide a reference-free representation of the phenomena under observation (Lachaux et al., 2003). A single bipolar montage (i.e., TB 1-2) is referred to as a “channel” throughout this study. Preliminary visual inspection of the iEEG recording and manual artifact rejection procedures excluded an average of 14 % of all trials (range: 8–22 %) with interictal activity across participants. This procedure decreased the risk of including trials modified by epileptic activity.

### 2.4. Channels and trials selection

Comparing subsequent causality estimates across subjects requires an equal number of channels and trials for each patient. We only included channels that do not share a common contact to avoid spurious increases in connectivity. There was a maximum of 30 channels that obeyed this rule for one patient. Reducing the channel numbers to 30 in all other patients required a further selection process. We manually selected channels localized in grey matter (based on MRI images) and visually recognizable neural responses to the stimuli. Furthermore, we included only the first 64 trials for each patient (minimal number of successful trials for the worst-performing subject). Therefore, with an a priori selection, we analyzed 18 subjects with 30 channels per subject, i.e., a total of 540 channels (Fig. 1a), 308 of which were in the left hemisphere and 232 in the right hemisphere.

**Figure 1:**
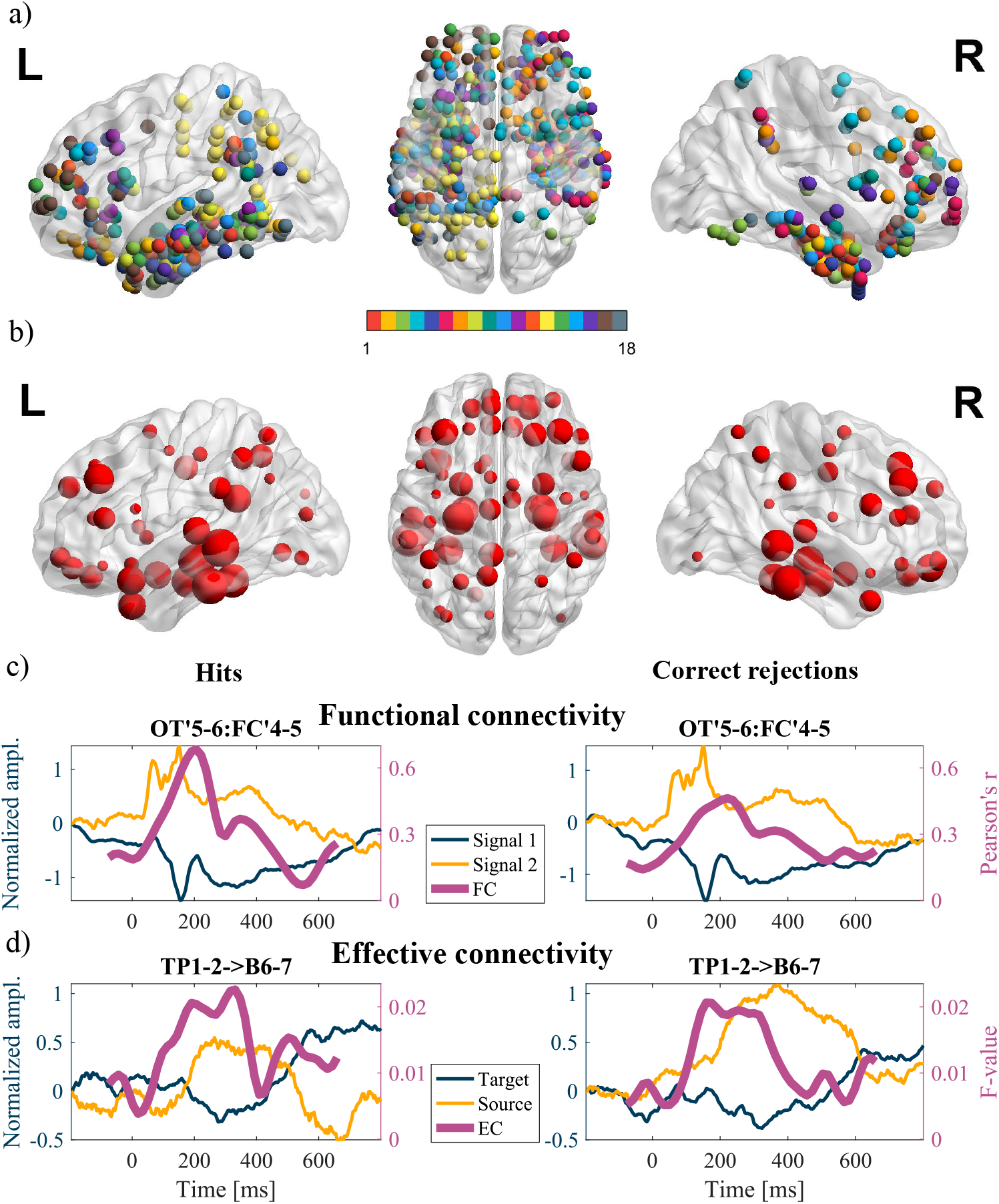
Recordings and connectivity. a) Overview of all recording locations across subjects. We recorded the brain activity of 18 epileptic subjects using multiple intracranial depth electrodes that targeted different brain regions. For each subject, we analyzed 30 bipolar channels, resulting in a total of 540 channels. The different colors corresponded to different subjects. b) We mapped these channels to the AAL atlas based on their MNI coordinates, covering 68 out of 90 possible regions with different channel densities. The size of each sphere corresponds to the sampling density of the region. c) Examples of dynamic correlation (i.e., functional connectivity) for a given channel pair (OT’5-6 and FC’4-5) for both Hits and Correct rejections. Note that a correlation is an undirected measure. d) Examples of dynamic Granger causality (i.e., effective connectivity) for a given channel pair (TP1-2 and B6-7) for both Hits and Correct rejections. According to the definition of Granger causality, the source influences the target.

### 2.5. iEEG preprocessing

iEEG preprocessing consisted of downsampling each channel to 256 Hz (original sampling frequency for two subjects) and subtracting the ensemble mean across trials to ensure stationarity (Barnett & Seth, 2014). We analyzed the 200 ms pre-stimulus baseline and 800 ms after the stimulus onset. To perform sliding-window connectivity analyses, we segmented each trial into windows of 64 samples (250 ms). We used a shift of 4 samples between two consecutive windows (similar results were obtained with a window of 32 samples). Each sliding-window was multiplied by a Hanning window to suppress spurious connectivity and reduce sensitivity to outliers (Preti et al., 2017). All data were processed with MATLAB (The Mathworks).

### 2.6. Connectivity analyses

We investigated functional (FC) and effective (EC) connectivity in sliding windows. We estimated dynamic FC and EC for each of the 18 subjects. To compare global levels of connectivity, we calculated the mean connectivity for each subject. Conversely, we pooled all connectivity estimates across subjects to analyze lateralization or directionality because the implantation varied significantly.

#### 2.6.1. Functional connectivity

We estimated FC between two channels as the mean Pearson’s correlation coefficient across all trials (Fig. 1c).

#### 2.6.2. Effective connectivity

We used dynamic multivariate Granger causality (MVGC) to estimate the EC between channels. It implements a statistical, predictive notion of causality whereby causes precede and help to predict their effects. Classical Granger causality from *Y* to *X* (the degree to which the past of *Y* helps predict *X*, over and above the degree to which *X* is predicted by its past) can be formally written as the log-likelihood ratio:

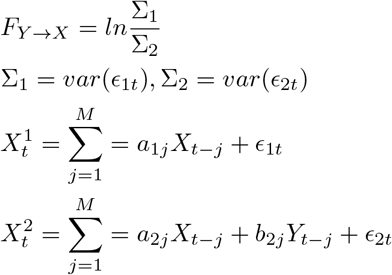

where *X* and *Y* represent recorded time series from two channels, *a* and *b* are parameters of the autoregressive process, *ɛ* represents residuals, and *M* is the model order (we used a constant order of 10, but similar results were observed using orders of 5 or 15).

We used the freely available toolbox from Barnett & Seth (2014) to calculate MVGC in overlapping sliding windows between all channels, separately for each patient. With the multivariate extension it is possible to control for common causal influences (Seth et al., 2015). Because our testing paradigm was time-constrained, and we used very short time windows, estimating the multivariate autoregressive (MVAR) model parameters might have been difficult. Nevertheless, we overcame this difficulty through the “vertical regression” implemented in the toolbox to address short time windows when multiple trials were available. This method is based on the assumption that each trial is an independent realization of the same underlying stochastic generative process. Therefore, we ended up with only one estimated MVAR model for all trials (Fig. 1d). We used the definition given by Gaillard et al. (2009), of feedforward direction as the causal influence of posterior channels onto the more anterior channel. If their *y* coordinates were identical, feedforward was defined as the causal influence of the lower onto the higher channel based on the *z* coordinates (this occurred in 3 % of the cases).

#### 2.6.3. Statistical testing

It is important to stress that in this study, we were limited by several factors such as the low number of subjects, short time windows, and tailored implantations, all of which are inherent to iEEG recordings. Moreover, the connectivity estimates followed a non-normal distribution. Therefore, for statistical testing, we used bootstrapping as a resampling technique, whereby random sampling with replacement from the distribution of interest is used to estimate the sampling distribution of almost any statistic (Efron & Tibshirani, 1994). Bias-corrected bootstrap was used in different ways according to the situation. This included calculating confidence intervals, testing differences between subjects, or testing increases compared to a baseline. Details of the procedures for statistical inference for each of these approaches are described in Appendix A.

### 2.7. Graph analyses

Two key concepts in graph theory are nodes and edges. In our analyses, nodes represent brain regions. We used the AAL atlas (Tzourio-Mazoyer et al., 2002) that parcellates the brain into 90 regions (including subcortical regions) to obtain identical parcellation for each subject. Each recording channel was assigned an area in the atlas based on its MNI coordinates using the SPM12 software package (Wellcome Department of Cognitive Neurology, London, UK) and the Anatomy toolbox (Eickhoff et al., 2005). Channels that did not belong to any region were not used in the mapping. Regions with no recorded signal were discarded, which resulted in the coverage of 68 out of the 90 AAL atlas regions (Fig. 1b). Note that in the AAL atlas, the perirhinal, parahippocampal, and entorhinal cortices are collectively referred to as the parahippocampal region.

The second constituent component of a graph is edges. We defined an edge between two regions as the mean MVGC of all corresponding channels. Since every graph can be represented as an adjacency matrix, and since we used a sliding window technique, our dynamic brain networks formed a set of adjacency matrices. Each adjacency matrix was based on data from all patients and represented an incomplete weighted directed graph. Traditionally, these matrices are thresholded and binarized to reduce measurement noise (Muldoon & Bassett, 2016), but arbitrary thresholding often leads to a loss of information (Rubinov & Sporns, 2011), and network measures are unstable across different thresholds (Garrison et al., 2015). Consequently, we opted to work with weighted directed graphs.

Two important concepts of network organization that might explain human cognitive abilities are segregation and integration. They provide essential insight into information processing and transmission. Segregation is the extent to which communication occurs primarily within tight-knit communities of regions. On the other hand, integration is the extent of communication between distinct regions. It is the ability of the network to integrate distributed information (Deco et al., 2015; Sporns, 2013). Both segregation and integration can be modeled with various measures (Bullmore & Sporns, 2009). We analyzed our dynamic memory networks in terms of efficiency and modularity (van den Heuvel & Sporns, 2013). All analyses were performed using The Brain Connectivity Toolbox designed for MATLAB (Rubinov & Sporns, 2010).

#### 2.7.1. Modularity

Modularity quantifies the degree to which the network may be subdivided into densely interconnected communities that maximize the number of within-group edges and minimize the number of between-group edges (Newman, 2006). We applied the iterative Louvain algorithm to the adjacency matrix with a resolution parameter of *γ* = 1 and random initial conditions for each time window of dynamic connectivity. A maximum of the modularity function across 10,000 runs was the resulting modularity with its accompanying network partition (Sporns & Betzel, 2016).

#### 2.7.2. Efficiency

Global efficiency is defined as the average inverse shortest path between any two nodes (Latora & Marchiori, 2001). Considering that it is linearly dependent on connectivity strength between nodes, we normalized it by dividing it by the mean connectivity across all non-zero edges.

#### 2.7.3. Null model

The use and choice of a null model are crucial in graph analyses (Hlinka & Hadrava, 2015; Hindriks et al., 2016). To create a stationary system with an identical covariance structure, we used an amplitude-adjusted multivariate extension of Fourier surrogates (MVFS) that matched the amplitude spectrum and amplitude distributions (see Schreiber & Schmitz 2000). We compared the magnitudes to the null models, i.e., we divided the obtained metric by a mean metric obtained in 1,000 surrogate networks (Westphal et al., 2017) and analyzed their dynamics. It is of note that Fourier surrogates could not be used to test the significance of global causality (Supp. Fig. 1). Moreover, we could not use the Erdos-Renyi null model due to incomplete brain coverage and the non-existence of certain links.

## 3. Results

Firstly, we compared global FC, approximated by Pearson’s correlation, for Hits and Correct rejections across all subjects and the entire brain. We observed a higher level of correlation for Hits across time (Fig. 2a, bootstrap *p* < 10^−16^). Unwrapping this analysis in terms of time showed that although the patterns were quite similar (Pearson’s *r* = 0.98, *p* < 10^−16^), there were significant differences between the two conditions starting at approximately 290 ms (bootstrap *p* < 0.05, FDR-corrected) (Fig. 2b). Because visual recognition memory relies more on the right than on the left hemisphere, we performed the same FC analyses focusing on each hemisphere. The patterns of left and right hemisphere FC were almost identical (Pearson’s *r* = 0.98, *p* < 10^−16^). The significant increase (bootstrap *p* < 0.05, FDR-corrected) in FC occurred slightly earlier in the right hemisphere (150 ms) than in the left (170 ms) (Fig. 2c).

**Figure 2:**
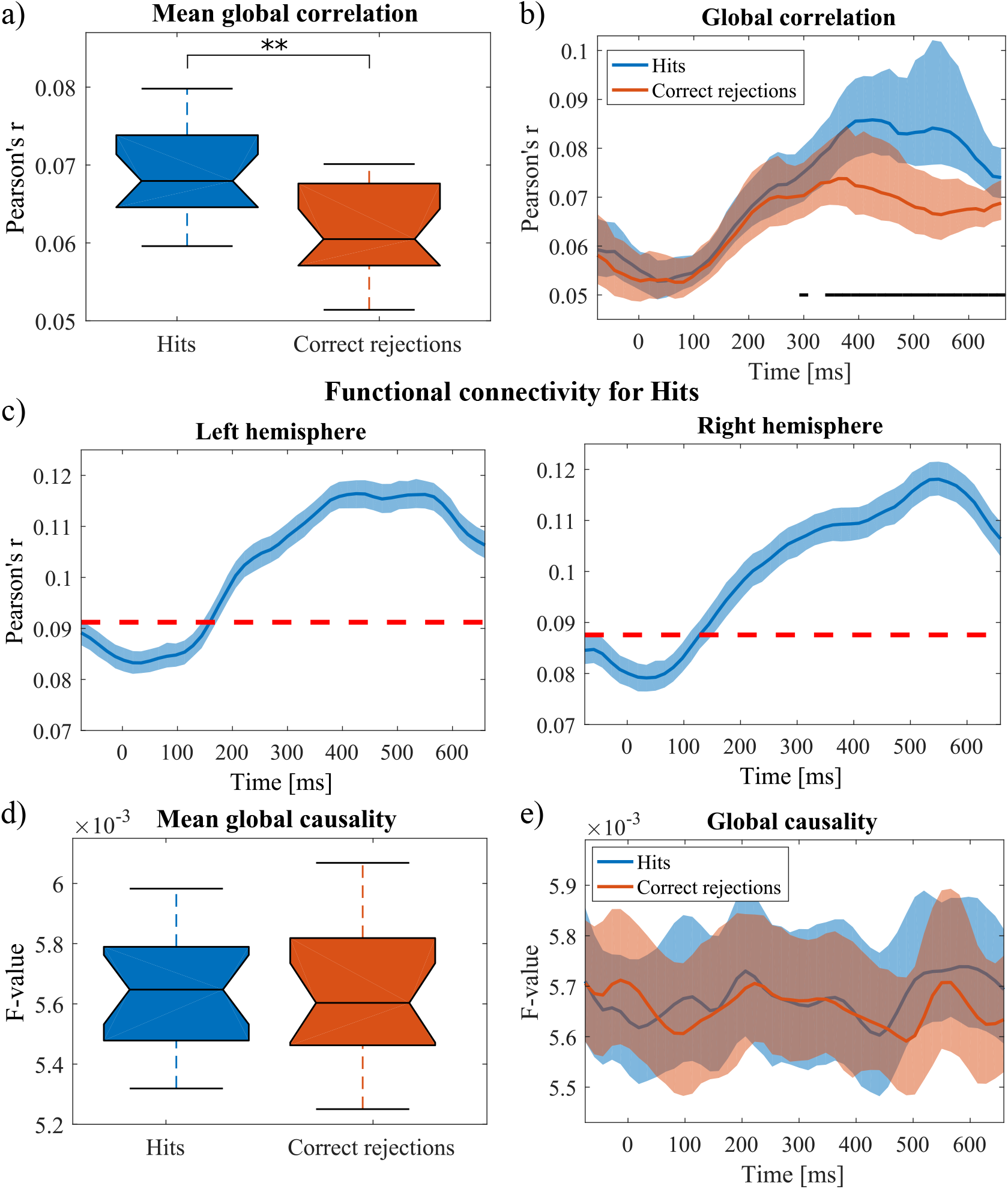
Connectivity analyses. a) We observed a higher level of global correlation for Hits than Correct rejections (*p* < 10^−16^). b) Resolved in time, there is a significant difference in correlation between the two conditions starting from 290 ms. The 90% bootstrap confidence interval is plotted in shaded colors. The black horizontal lines indicate significant time intervals. c) If we focus on the FC within each hemisphere, very similar temporal patterns can be observed (Pearson’s *r* = 0.98, *p* < 10^−16^). Compared to the baseline, we see a significant increase in both right hemisphere (starting from 150 ms) and left hemisphere correlations (starting from 170 ms). The red dotted lines represent the threshold for a significant change from the baseline. d) Unlike for correlation, the mean global causality for Hits was not statistically higher than for Correct rejection (*p* = 0.28). e) Moreover, the time courses of mean causalities were not significantly correlated (Pearson’s *r* = 0.22, *p* = 0.13), and there were no statistical differences in time (*p* > 0.05, FDR-corrected).

Since directionality cannot be tested by correlations, we further analyzed effective connectivity by Granger causality. We calculated MVGC using vertical regression in short sliding windows, thereby providing a dynamic estimate of causality strength. Averaged in time, MVGC for Hits and Correct rejections across subjects and the entire brain were not significantly different (bootstrap *p* = 0.28) (Fig. 2d). Although, the conditions showed different dynamics (Pearson’s *r* = 0.22, *p* = 0.13) no significant difference was noted in terms of time (bootstrap *p* > 0.05, FDR-corrected) (Fig. 2e).

Following these preliminary analyses, we analyzed dynamic causality in more detail by focusing on feedforward and feedback directionality within each hemisphere. We calculated partial Pearson’s correlations (controlling for global causality) between the time courses of these conditions. We found significant similarities between feedforward and right hemisphere causality time courses (*r* = 0.61, *p* < 10^−5^) (Fig. 3a). On the other hand, the feedback causality time course was closely connected with the left hemisphere (*r* = 0.70, *p* < 10^−7^) and across both hemispheres time course (*r* = 0.48, *p* < 10^−3^). It is importance to note that such significant correlations were not found for Correct rejection (Supp. Fig. 2), neither were they randomly obtainable. This was confirmed by shuffling the labels of the links (Supp. Fig. 3).

**Figure 3:**
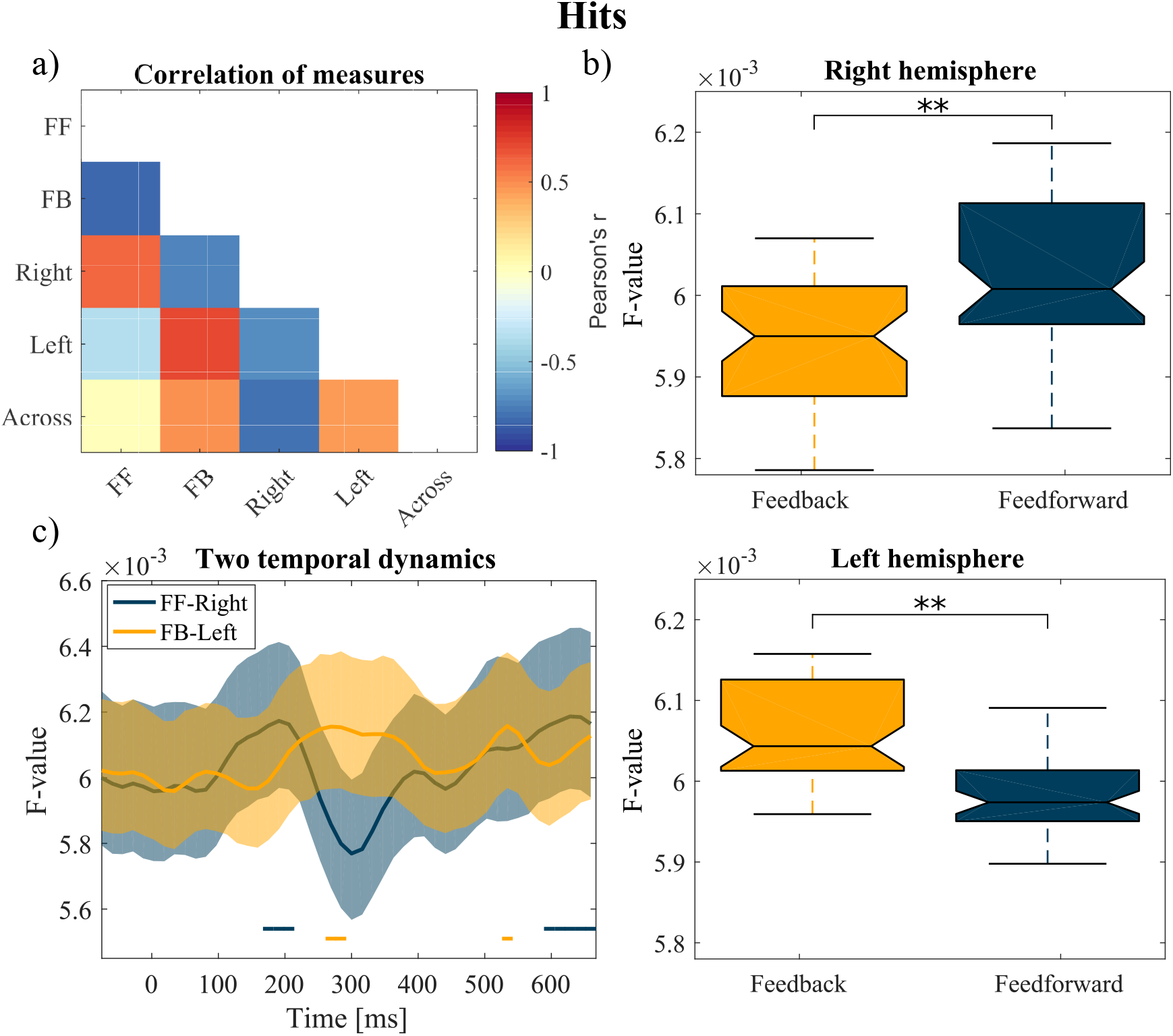
Effective connectivity analyses for Hits. a) We investigated the temporal evolution of causality based on directionality and lateralization. We found striking similarities for some of the patterns: right hemisphere and feedforward causality were highly correlated, as well as left hemisphere (or across-hemisphere) and feedback causality. These two modes were highly anticorrelated. Non-significant correlations (FDR-corrected) were set at 0. b) When time is averaged, we observe more feedforward causality in the right hemisphere (*p* = 0.005) and more feedback causality in the left hemisphere (*p* = 0.001). c) The right feedforward causality significantly increased as of approximately 170 ms and then decreased at 300 ms. This decrease was associated with a significant increase in left feedback causality. The 90% bootstrap confidence interval is plotted in shaded colors. Horizontal lines indicate periods of significant increase.

Further analyses of Hits showed that the causality of feedforward connections was higher than that of feedback connections in the right hemisphere (bootstrap *p* = 0.005), while the reverse was true in the left hemisphere (bootstrap *p* = 0.001) (Fig. 3b). Furthermore, the significant increase in the right hemispheric feedforward causality occurred much earlier (170 ms, bootstrap *p* < 0.05) than the left hemisphere feedback causality (270 ms, bootstrap *p* < 0.05) (Fig. 3c).

To further improve our understanding of the networks supporting visual recognition memory, we switched to data-driven analyses (i.e., whole-brain rather than by hemisphere). Therefore, we focused on two metrics that describe network topology: modularity (a measure of segregation) and efficiency (a measure of integration). Network topology changed over time (Fig. 4a). We observed a highly segregated (modular) topology from 110 ms after stimulus onset. It then transitioned into a more integrated (efficient) topology at approximately 220 ms. Moreover, the significant increase in modularity (MVFS *p* < 0.05, FDR-corrected) occurred just before the increase in right hemisphere feedforward causality (Fig. 4b). Similarly, the increase in efficiency occurred just before the increase in left hemisphere feedback causality (Fig. 4c). These results suggest that changes in network topology could precede (and maybe drive) changes in information flow. We observed similar modularity and efficiency patterns even with a different brain parcellation, namely the Harvard-Oxford atlas (Pearson’s r between dynamic modularity from AAL and Harvard-Oxford atlas *r* = 0.51, *p* < 10^−3^, resp. efficiency *r* = 0.47, *p* < 10^−3^).

**Figure 4:**
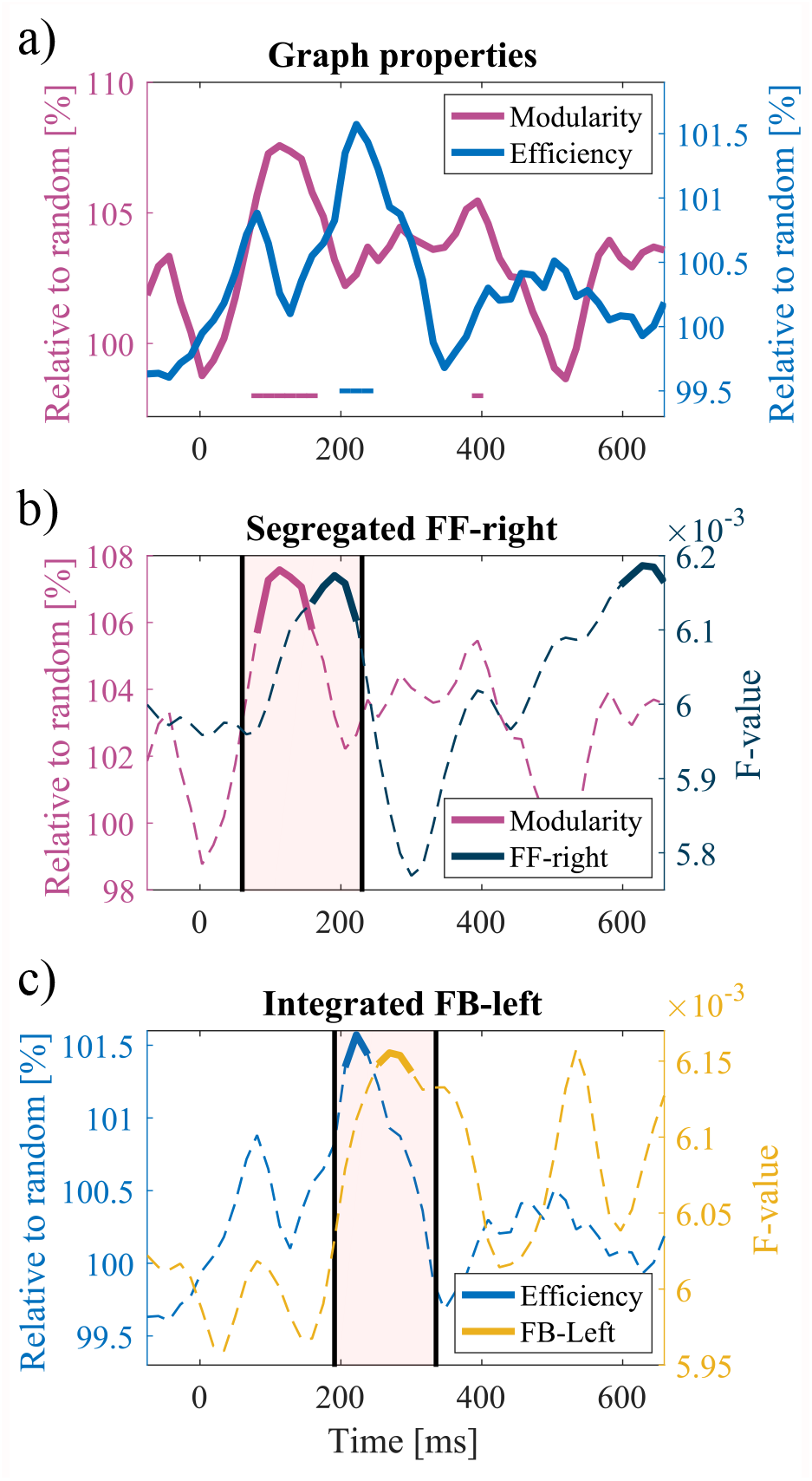
Changes in the network topology across time. a) At approximately 110 ms, the network shows a more modular topology. This segregated state is followed by a more integrated structure characterized by higher efficiency at 220 ms. Horizontal lines represent periods of significant increase/decrease. b) After the first peak of modularity at 110 ms, a significant increase in right hemisphere feedforward causality at approximately 150 ms can be observed. c) In addition, the first peak of efficiency at 220 ms precedes a significant increase in left hemisphere feedback causality at 250 ms. Solid lines indicate significant values. The rectangles highlight the time intervals of interest.

In addition, we used the Louvain algorithm to detect community structures in networks. We consistently identified three main communities at each time window, i.e., three highly interconnected subgraphs (Fig. 5a). Based on the most frequent allegiance of each node, the first community comprised regions of the right temporal lobe as well as many frontal regions bilaterally. The second community comprised regions in the left hemisphere, mostly from the temporal lobe and other parietal and frontal lobes. Interestingly, both the left and the right hippocampi were more linked to this second community. The third community comprised the left parahippocampal and inferior frontal gyri as well as the left amygdala.

**Figure 5:**
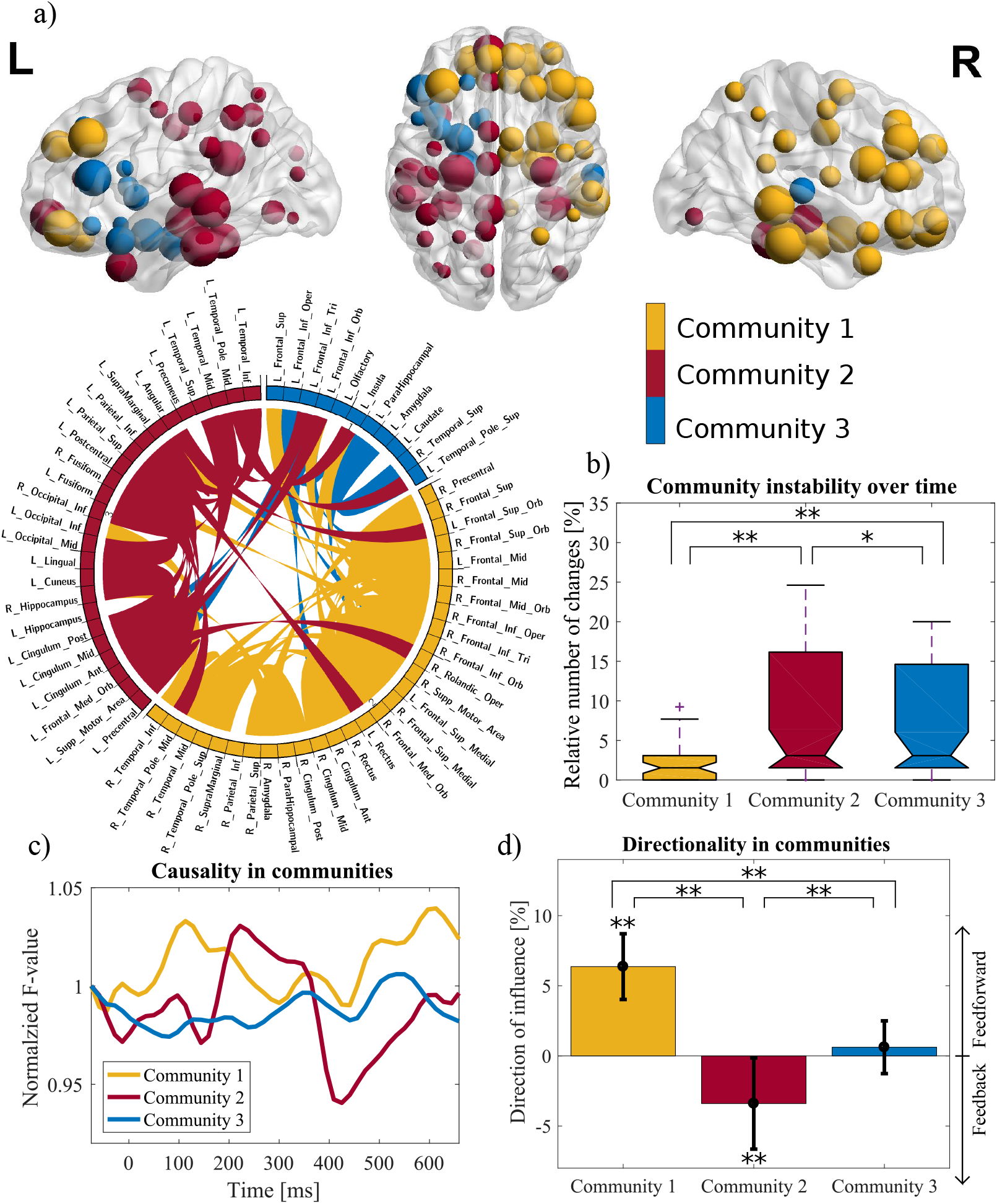
Network communities. a) The Louvain algorithm for community detection consistently identified three main communities. The first community comprises regions of the right visual stream and medial temporal lobe structures as well as frontal regions in both hemispheres. The second community comprises regions in the left MTL and the right hippocampus. The third community comprises the left parahippocampal gyrus, left amygdala, and left inferior frontal gyrus. The size of the spheres in the brain graph corresponds to the nodal strength. For representation purposes, the circular form (Krzywinski et al., 2009) shows only the 3 % consisting of the strongest links. b) Even though the core of each community remained stable across the time course, some nodes changed allegiance. The first network shows the highest stability, i.e., the lowest instability index defined as a relative number of allegiance changes per node. c) The first community showed the earliest increase in causality (110 ms). The second community followed at approximately 220 ms. d) Temporal average standard deviation of the direction of influence is significantly different between communities. Moreover, the first community exhibits a significant prevalence of feedforward causality. Conversely, the second community displays a significant prevalence of feedback causality (* *p* < 0.05, ** *p* < 0.001, FDR-corrected).

**Table 1:** The relation between connectivity and performance. We correlated the typicality of the subjects’ right feedforward and left feedback connectivity with their performance on recognition memory tasks as assessed by minimal reaction time (an index of the speed to perform the task) and d-prime (an index of the ability of the subjects to discriminate Hits from Correct rejection). Results are comparable for both functional and effective connectivity analyses. We obtained identical p-values using permutation testing with 10,000 repetitions.

|  | Minimal Reaction Time |  | d-prime |  |
| --- | --- | --- | --- | --- |
|  | Functional | Effective | Functional | Effective |
|  | Pearson's r (p-value) | Pearson's r (p-value) | Pearson's r (p-value) | Pearson's r (p-value) |
| FF-Right typicality | -0.34 (0.10) | -0.32 (0.13) | 0.04 (0.44) | 0.15 (0.16) |
| FB-Left typicality | 0.26 (0.16) | 0.59 (0.006) | -0.52 (0.98) | 0.06 (0.43) |

Some nodes changed their allegiance throughout the time course, but the core of each community remained stable (Supp. Fig. 4). The communities differed in their stability (one-way ANOVA, *p* < 10^−3^), with the first community being the most stable, i.e., having the lowest instability index defined as the relative number of node allegiance changes over the course of time (Fig. 5b). Therefore, if a node changed allegiance, it was mostly between the second and third community.

The first community showed the earliest increase in causality compared to baseline, at a similar timing as the increase of feedforward causality in the right hemisphere (Fig. 5c). Furthermore, the first community feedforward causality correlated significantly with that of the right hemisphere identified in the first set of analyses (*r* = 0.76, *p* < 10^−9^). Likewise, the causality increase in the second community occurred later, in approximately 230 ms (Fig. 5c). The feedback causality correlated significantly with the feedback causality in the left hemisphere (*r* = 0.41, *p* = 0.004). The causality of the third community remained comparable to the baseline. Moreover, in the analysis of the directionality of influence, defined as the ratio between feedforward and feedback causality, the communities differed significantly (one-way ANOVA, *p* < 10^−16^). The first community exhibited a significant prevalence of feedforward interactions (bootstrap *p* < 0.05, FDR-corrected), while the second showed a prevalence of feedback interactions (Fig. 5d).

All in all, the two functional systems identified on the basis of a hypothesis regarding the hemispheric lateralization of visual recognition memory are also identifiable with a data-driven community analysis. However, the community analysis offered a more detailed delineation of participating structures than a simple dichotomy between the right and left hemispheres.

Finally, we associated the observed connectivity patterns with the subjects’ performances. We assumed that a higher performance level was associated with neural activity that resembles typical activity. Moreover, we expected the feedforward causality to drive fast response, unlike the feedback. Therefore, we calculated a one-sided Pearson correlation between the typicality of neural response (correlation between subjects’ causality time course and the template of right feedforward and left feedback causality - from Fig. 2c for FC and from Fig. 3c for EC) and the minimal reaction time and d-prime (Tab. 1). Although mostly non-significant, the descriptive analysis showed that the right hemisphere feedforward causality were negatively correlated with minimal reaction times (the more typical the neural response, the faster the minimal reaction times; *r_FC_* = − 0.34, *p* = 0.10; *r_EC_* = − 0.32, *p* = 0.13) while the left hemisphere feedback causality showed a positive correlation (*r_FC_* = 0.26, *p* = 0.16; *r_EC_* = 0.59, *p* = 0.006) (Supp. Fig. 5). In terms of d-prime, we tested for positive correlation and did not find consistent results across measures. We obtained identical p-values using permutation testing with 10,000 repetitions.

## 4. Discussion

A typical characteristic of visual recognition memory is its rapidness. Subjects in this study were able to respond correctly in less than 600 ms, with the fastest correct responses being approximately 370 ms, which is consistent with previous results (Besson et al., 2012). Although the perirhinal cortex and hippocampus have been identified as the core brain regions that support recognition memory, studies have consistently reported many other brain regions, mainly in the temporal lobe, but also in the frontal and parietal lobes (Gonzalez et al., 2015; Hoppstädter et al., 2015; Rutishauser et al., 2018; Despouy et al., 2020). Nevertheless, little is known about the relationships between all these brain regions and how these relationships evolve over time. The amount of functional networks that are activated during recognition memory is also unknown. Therefore, there is a need to understand the dynamical organization of the large-scale functional brain networks that underlie recognition memory on a millisecond-by-millisecond scale (Kopell et al., 2014; Sporns, 2014).

Because this endeavor requires a high spatial and temporal resolution, we analyzed functional and effective connectivity of intracerebral EEG in short sliding time windows to track connectivity changes and information flow during a visual recognition memory task. We identified large-scale brain networks involved in successful recognition. The first network mainly involved the right visual ventral stream and the bilateral frontal regions. It was characterized by predominant feedforward activity, starting rapidly in approximately 110 ms post-stimulus and peaking at 190 ms, modular topology, and high stability. It was followed by the involvement of a second network, predominantly in the left hemisphere, but notably also involving the right hippocampus, characterized by predominant feedback activity which peaked at 270 ms, integrated topology, and lower stability. It is important to note that the patterns of right hemisphere / feedforward and left hemisphere / feedback connectivity were found only for Hits, but not for Correct rejections. Interestingly, the peaks in modularity and efficiency (the transitions from less segregated to more segregated and from less integrated to more integrated topology) preceded the peaks in right / feedforward and left / feedback connectivity, which suggest a causal link between changes in network topology and modes of information processing. A third, lower-scale network, was identified and was related to the second. Overall, these results confirm that several large-scale brain networks, each with specific properties and dynamics, rapidly unfold (i.e., in less than 300 milliseconds) during recognition memory. These networks involve many brain regions bilaterally, even for such a basic cognitive capacity.

### 4.1. Different networks unfold rapidly in time

We performed two types of analyses to identify the networks that support recognition memory. The first one was driven by the hypothesis that there would be a high level of asymmetry between both hemispheres. The second was data-driven (it involved all recorded brain regions with no a priori selection of brain regions or hemispheres) and was based on the identification of brain region communities (by maximizing the number of within-group edges and minimizing the number of between-group edges (Newman, 2006)). Both analyses were carried out dynamically over the entire time period. It is important to note that both were convergent, demonstrating robust findings, although the data-driven analysis provided complementary information.

The hemispheric analyses revealed a robust functional difference between the right and left hemisphere. The right was mainly characterized by a feedforward information flow, while the left mainly by the feedback information flow. The difference between the amount of feedforward and feedback connections was significant within each hemisphere (Fig. 3b). Interestingly, the dynamics of the two hemispheres were different since the peak of the feedforward information flow in the right hemisphere occurred in approximately 170 ms. In contrast, the peak of the feedback information flow in the left hemisphere occurred later, in approximately 270 ms, at a moment when the feedforward information flow in the right hemisphere sharply decreased (Fig. 3c).

The data-driven analysis identified three networks. The first encompassed many brain regions in the right temporal lobe and the bilateral frontal lobes. Interestingly, this network showed a very rapid increase (between 100 and 200 ms) in effective connectivity. It was characterized by effective predominant feedforward connectivity, which is consistent with the facts already known about the rapidity and flow of information from the visual ventral stream (VanRullen & Thorpe, 2001; DiCarlo et al., 2012). The second encompassed brain regions in the left temporal lobes, as well as the parietal and frontal lobes. It is highly important to note that it encompassed both the left and the right hippocampi. The effective connectivity of this second network peaked later than the first, albeit rapidly after stimulus onset between 200 and 300 ms. It was characterized by effective predominant feedback connectivity. A third network comprised regions in the posterior frontal and anterior temporal lobes. Unlike the two previous networks, it did not have a clear information flow direction. The first network was characterized by high community stability (few nodes changed allegiance over the periods). The second and third communities were less stable, with nodes interchanging allegiance throughout the period. Follow-up analyses showed that the first connectivity network pattern was very similar to the feedforward connectivity pattern observed in the right hemisphere. Likewise, the second network connectivity pattern was very similar to the feedback connectivity pattern observed in the left hemisphere. Overall, these analyses provide the picture of three functional networks that underlie visual recognition memory, each with specific topography, temporal dynamics, preferred direction of information flow, and stability. In other words, within 300 ms, the brain undergoes a massive dynamic functional reorganization phase that involves several networks.

### 4.2. A large-scale network account of recognition memory

The first functional network was partly expected since previous intracerebral EEG studies had already demonstrated a high early involvement of the right visual ventral pathway in visual recognition memory (Barbeau et al., 2008; Despouy et al., 2020). Furthermore, it has already been established that frontal lobe regions are involved in recognition memory (Swick & Knight, 1999; Bastin et al., 2006; Despouy et al., 2020), as early as 110 ms after stimulus presentation (Bar et al., 2006; Barbeau et al., 2008). The identification of a second network was more unexpected, particularly a network that tended to be more left-sided and prominently characterized by feedback connectivity.

Although further studies will be required to clarify each network’s role, they highly correspond with current knowledge of the neurocognitive architecture that underlies recognition memory. Familiarity and recollection are the two processes that underlie successful recognition memory (Brown & Aggleton, 2001). Familiarity is a fast process that relies mainly on the perirhinal cortex as the core brain region, along with the ventral visual pathway. It does not include the hippocampus (Eichenbaum et al., 2007). Therefore, the first network could be mainly involved in familiarity. In contrast, recollection is assumed to be a slower process that relies on the hippocampus as the core brain region and the extended hippocampal system in general, which involves relays in the mammillary bodies, anterior thalamus, cingulate cortex, and parietal lobes (Yonelinas et al., 2005). Therefore, the second network could be more involved in recollection, as suggested by the fact that it was delayed compared to the first network, but also by the fact that both right and hippocampi belonged to this network.

While familiarity depends mainly on processing the world surrounding the subject (i.e., bottom-up processes), recollection on the other hand, requires interactions with the internal world (i.e., memory) to retrieve the spatio-temporal context of occurrence of the stimuli that needs to be recognized. The notion that the first network is characterized mainly by feedforward connectivity while the second is characterized by feedback connectivity is consistent with these hypotheses. Recently, Kar et al. (2019) showed the advantage of recurrent computations for object recognition. Increased feedback connectivity could represent top-down modulation (Ishai et al., 2006), the build-up of an internal representation of the stimulus (Rijsbergen & Schyns, 2009), or access to a distributed semantic system (Burke et al., 2014). This is supported by the notion that the second slower nework could be related to language or the need for internal speech, which would explain why it is more left-lateralized.

Recently, Bastin et al. (2019) proposed a large-scale functional architecture that supports familiarity and recollection. This Integrative Memory Model emphasizes the large number of brain regions involved in familiarity and recollection processes. Moreover, it proposed that an “attribution and attention” system, mainly dependent on frontal lobe regions, was involved in recognition memory. This system involves top-down attention, activity maintenance, metacognitive knowledge, and monitoring and decision-making, leading to the subjective feelings and explicit judgments that occur during recognition memory. The fact that the first and third networks encompassed many frontal lobe regions is consistent with this proposal. It may also explain why the second and third networks are less stable than the first and exchange node allegiances over time if monitoring and decision-making are underway.

A switch from a goal-oriented network (familiarity) to an introspective one (recollection) requires significant reorganization of the brain, which also involves the hippocampus (Barbeau et al., 2017). Previous studies have independently identified network changes occurring after 240 ms during recognition memory tasks (Barbeau et al., 2008; Maillard et al., 2011), which lends support to the idea that this switch between networks occurs. Studies that focus on functional connectivity using fMRI have consistently revealed brain network reorganizations during cognitive tasks (Bola & Borchardt, 2016; Ekman et al., 2012; Shine et al., 2016). Note that the switch (between external and internal worlds) also involves the frontal lobes (Brincat & Miller, 2016). Westphal et al. (2017) suggested that a cross-talk between two large-scale networks during episodic memory may push the brain into a globally more integrated state, enabling higher information transfer fluidity. This increase in global integration is driven by an increase in cognitive load, whereby the brain can adopt a more global workspace configuration (Kitzbichler et al., 2011; Ekman et al., 2012). Moreover, higher task demands were already noted to decrease modularity (Vatansever et al., 2015). Overall, network topology is tightly linked to information transmission (Lynn et al., 2020). These notions are consistent with our findings of a critical switch between different networks that precede a change in information flow, and underscore the brain’s ability to reconfigure dynamic networks in response to changing cognitive demands (Cohen & D’Esposito, 2016).

It is highly noteworthy that these findings indicate that large-scale functional networks can have several modes of relationships, for instance, critical moments of transition between networks (such as between the first and second network) or strong interactions (such as between the second and third network where node allegiance fluctuates between the two networks over time). Overall, this study provides a richer and more integrated picture of the brain networks that underlie recognition memory.

### 4.3. Recognizing stimuli: Hits vs. Correct rejection

It is of note that the pattern of predominantly feedforward and feedback information flow observed in the right and left hemispheres was identified only for Hits but not for Correct rejections. The visual recognition memory task used in this study was based on a go/no-go paradigm. The response (raising fingers from a response pad) was provided only for Hits, while CR did not require a response. This paradigm was chosen because it forces subjects to use their fastest strategy (Barragan-Jason et al., 2013). Consequently, Hits required the involvement of more brain regions than CR.

Watrous et al. (2013) suggested that functional connectivity related to correct versus incorrect context retrieval was rather global than regionally specific. We consistently found significant differences between Hits and CR in global FC from approximately 290 ms. These increased functional interactions are believed to be a signature of successful recollection (Schedlbauer et al., 2014; King et al., 2015).

If findings are not related to behavioral performance, there is a risk that they may reflect non-psychological factors. Therefore, we tried to verify whether the right feedforward connectivity pattern, possibly underlying familiarity, drove fast behavioral responses. The left feedback pattern, possibly related to recollection, was associated with slower responses (Serre et al., 2007; Yoo et al., 2019). We found consistent results supporting this idea. To be specific, both functional and effective feedforward connectivity were negatively correlated to reaction times, while positive correlations were found for feedback connectivity. However, these correlations did not have a high statistical power. The number of subjects or the variability of the implantations (impacting the typicality of the neural responses) may have decreased the statistical power. Even though Shine et al. (2016) suggested a direct link between cognitive performance and dynamical brain network reorganization, we found no significant correlation between modularity or efficiency and performance, probably due to the factors just mentioned.

### 4.4. Challenges of whole-brain, dynamic, connectivity analyses using iEEG

Intracerebral EEG has the tremendous advantage of providing an excellent spatial and temporal resolution not provided by other methods. However, it also has drawbacks that may have impeded the connectivity analyses. Very few studies focus on whole-brain, dynamic, effective connectivity using intracerebral EEG data because of the specific challenges posed by this approach. Most of the previous studies predefined regions of interest a priori (Staresina et al., 2012) and did not focus on large-scale networks or their temporal dynamics (see exceptions such as Gaillard et al. 2009). In fact, iEEG analyses pose specific challenges such as the relatively low number of subjects, short and non-stationary time-series, and tailored electrode implantations, which may under-sample some brain regions. Moreover, the large number of sampled brain regions requires synthesizing information across edges and nodes. Some patients also participated in more trials than others; however, we had to restrict our analyses to the same number of trials per patient because the number of trials directly influences the magnitude of connectivity estimates.

To overcome these issues, we pooled results from all 18 patients and mapped channel locations to the AAL atlas. We were thereby able to reconstruct signals from 68 out of 90 brain regions. Using vertical regression in all trials (limited to 64 per patient), we could estimate causality in short (250 ms) and stationary time windows. Dedicated statistical analyses had to be designed at each stage of the analyses to assess the value of the findings. In addition, we followed the definition by Gaillard et al. (2009), of feedforward and feedback processes, but this was only a rough simplification. In contrast, a hierarchical anatomically based model might be better to represent brain processes (Markov et al., 2014). Furthermore, functional rather than anatomical parcellation could be better to associate network topology and behavioral responses (Salehi et al., 2020). Future studies could also benefit from frequency-resolved measures to detect networks that operate on specific frequencies (Mao et al., 2016). Overcoming these obstacles could open new paths for future studies on connectivity dynamics using iEEG.

It is worth mentioning that iEEG involved recordings from epileptic patients. Therefore, epilepsy could impact the generalization of the results. However, as in all similar studies, we removed the interictal activity periods recorded simultaneously with the task. Previous studies have also shown that similar ERPs, characterized by latency, morphology, and amplitudes, are found across independent studies and epilepsy centers (e.g., Trautner et al. 2004 vs. Barbeau et al. 2008). Interestingly, a recent study combining iEEG and fMRI demonstrated only small functional neuroanatomical differences during an episodic memory task between a group of epileptic patients and a group of matched healthy subjects (Hill et al., 2020). Overall, despite the limitations, iEEG studies appear to provide useful and reliable information.

## 5. Conclusion

In conclusion, this study reveals novel findings regarding the dynamics of the large-scale functional networks that underlie recognition memory. It could generate hypotheses that could be specifically tested in future work. For example, during a recognition memory task, neuronal activity should mainly reflect feedforward and early activity in the first right hemisphere network (Sugase et al., 1999). Such activity should also differ from the neuronal activity recorded in the second network. It would also be interesting to examine the physiological mechanisms that enable the transition between different networks. In general, this study shows that whole-brain dynamic connectivity analyses using intracerebral EEG offer a promising avenue to study different classes of cognitive abilities.

## Acknowledgement

This work has been supported by Campus France grant No. 44815QG, the Czech Science Foundation project No. 19-11753S, project No. LO1611 with financial support from the MEYS under the NPU I program and from the Specific university research – grant No. A2 FCHI 2018 012.

## Data availability statement

The data may contain clinical information and cannot be shared publicly.

## Appendix A. Bootstrap statistics

This is a description of the different methods that we adapted and designed to test the statistical significance of differences between conditions and increases in connectivity, or to calculate confidence intervals. We used a resampling method called bootstrapping. It uses random sampling with replacement from the distribution of interest to estimate the sampling distribution of almost any statistic (Efron & Tibshirani, 1994). To be more specific, we used the bias-corrected implantation that corrects for bias and skewness in the distribution of bootstrap estimates. For a detailed description, see Penn (2020).

We calculated confidence intervals of dynamic connectivity (Fig. 2b) by creating a bias-corrected bootstrap distribution of mean connectivity values at each time-point. We used the hybrid method where a 90% confidence interval from 10,000 repetitions is plotted around a mean value of the original distribution.

We tested whether there was a difference in mean value between two conditions across subjects, i.e., whether the mean connectivity of Hits was significantly different from the mean connectivity of Correct rejections (as in Fig 2a,b,d,e), using a two-tailed paired sample bootstrap test with 10,000 repetitions. The null hypothesis for the test was that the mean connectivity calculated for the difference between Hits and Correct rejections was equal to zero. In Fig. 2b,e, the resulting p-values were corrected for multiple comparisons in the time domain with the FDR algorithm to take into account the 48 tests performed across time. In Fig. 5c, we used the same approach to test whether the mean direction of influence was zero in each community

The original bootstrap method is designed for independent identically distributed data. A standard bootstrap is not appropriate when data samples are dependent (such as time series). Therefore, we used a stationary bootstrap - a block technique that attempts to preserve the underlying autocorrelation (Lahiri, 2003). TThis technique is based on a circular wrap of data (end-to-start wrap around the data around a circle) and a random window length that removes the edge effect of uneven weighting at the beginning and the end (Politis & White, 2004). To test whether there was a significant difference in mean feedforward connectivity between the left and right hemispheres (Fig. 3b), we compared the two corresponding stationary bootstrap distributions and calculated a p-value, as mentioned above. Finally, to test whether there was a significant increase in a time course (as in Fig. 2c or Fig. 3c), we created a bias-corrected bootstrap distribution of mean connectivity in each time window by randomly sampling subjects with repetitions 10,000 times. Furthermore, we compared each bootstrap distribution to the baseline bootstrap distribution (from a time window centered at −75 ms) to obtain the resulting p-values.

---

**Supplementary Figure 1:**
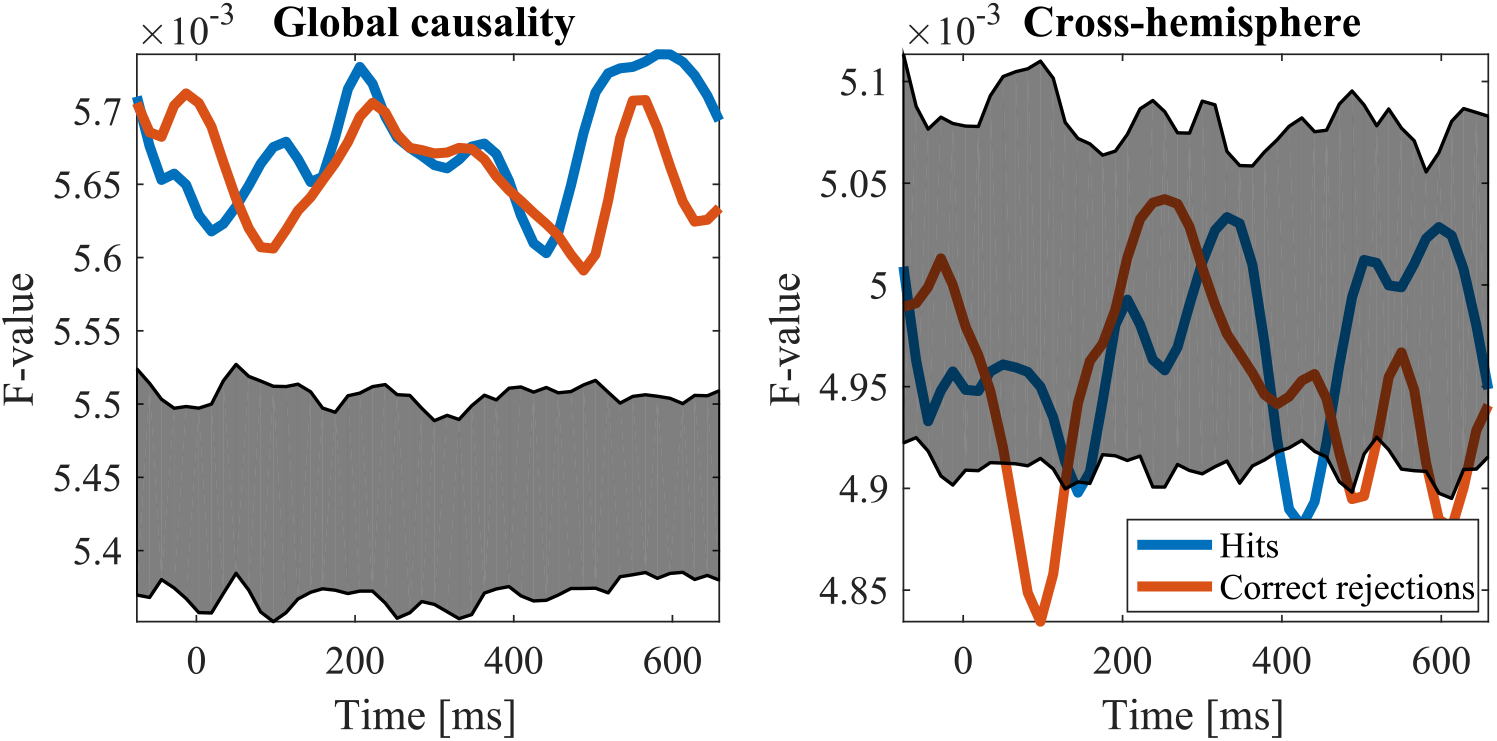
The significance of the causality time series could be assessed by comparison to corresponding surrogates. Nevertheless, in our analysis, all time series (except across hemispheric causality) have values above the 90 % confidence interval of amplitude-adjusted multivariate extension of Fourier surrogates. This is why we do not focus on the examination of across hemisphere causality. While the surrogates correspond well to individual pairs, they cannot be used to assess the significance of mean connectivity across all pairs. It might be due to the non-stationarity of the underlying process and the stationarity of surrogates. It would appear that in every time point, there are highly significant causality values that drive the mean value to be significant as well.

**Supplementary Figure 2:**
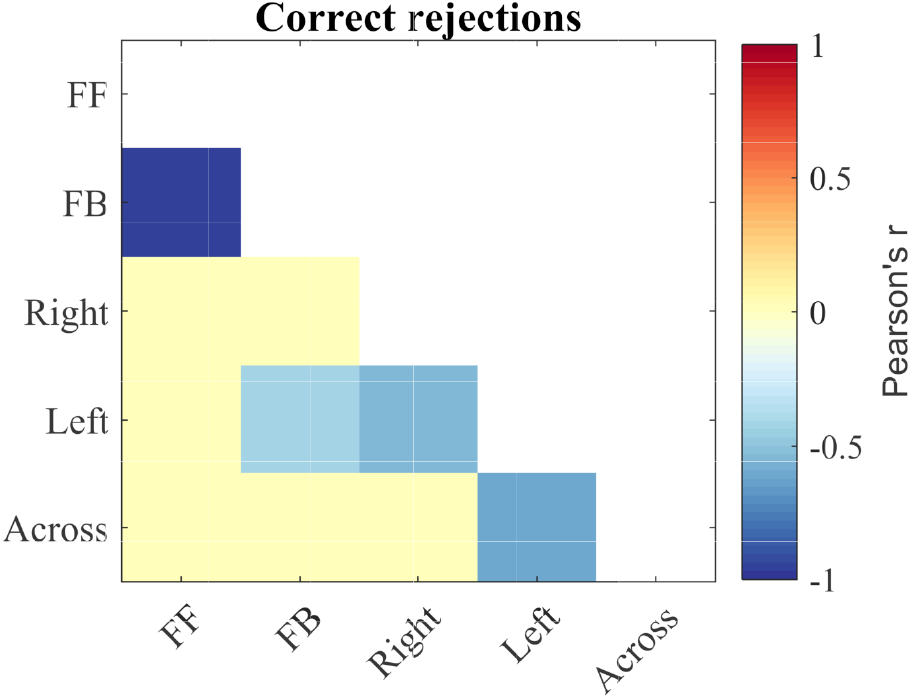
In Hits, we found significant similarities between causality time series. Therefore, we calculated the partial Pearson’s correlation (controlling for the effect of global causality) between different causality time courses for Correct rejections. We did not observe a significant positive correlation for directionality and lateralization, as was the case in the main analysis.

**Supplementary Figure 3:**
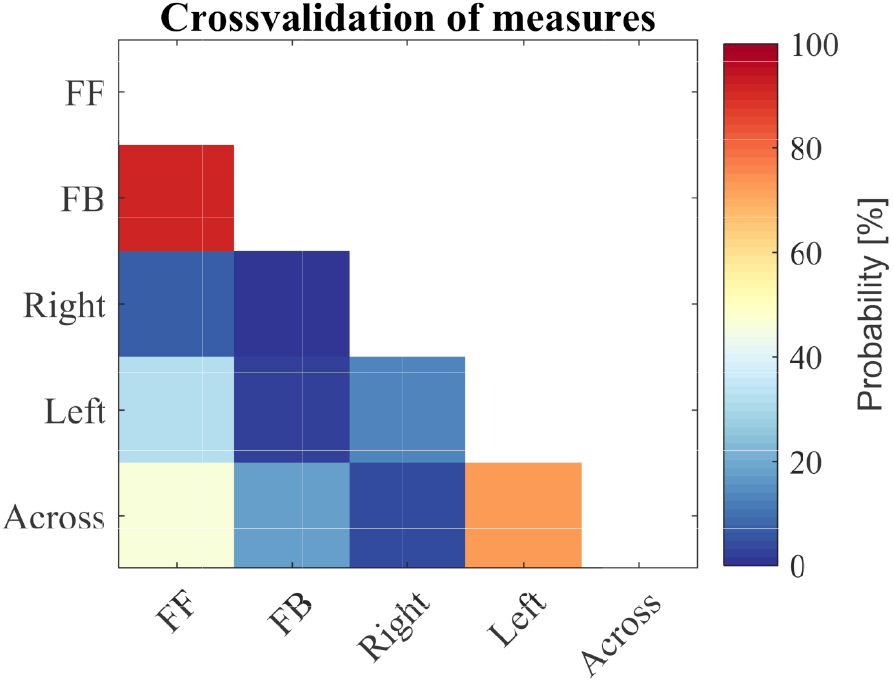
We tested whether the correlations obtained in 2a were randomly obtainable. We randomly shuffled indices regardless of whether a link was feedback/feedforward, cross-hemispheric, or within the left/right hemisphere. The number of links in each category remained the same as in the original analysis. Moreover, one link cannot feedforward and feedback simultaneously (same for other classes). We repeated this procedure 10,000 times. We plotted the probability of obtaining a higher correlation coefficient than in the main analysis (higher for positive correlations, reps. lower for negative correlations). The probability of obtaining higher magnitudes of right feedforward and left feedback correlations is less than 5%.

**Supplementary Figure 4:**
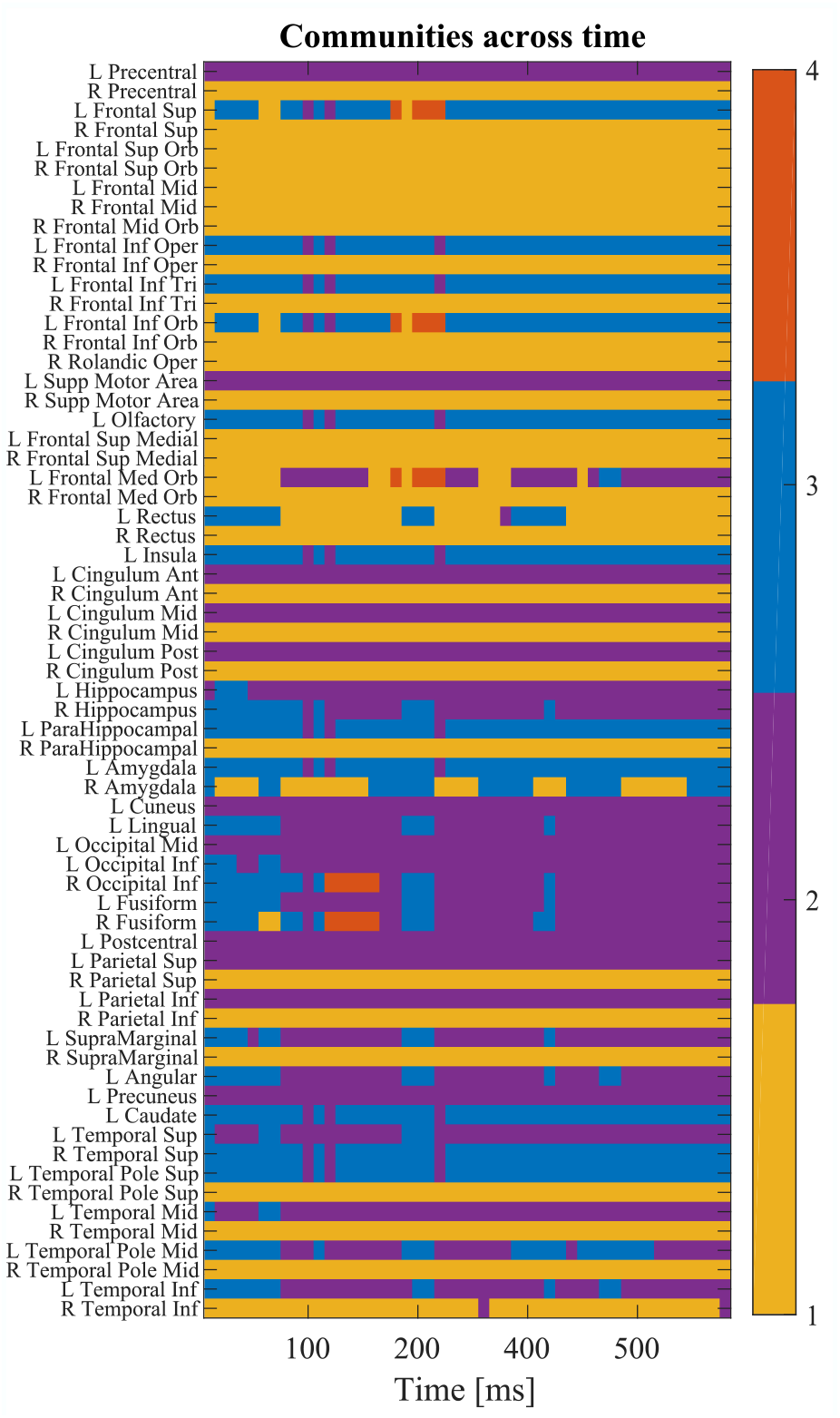
In each time window, we assigned a community index to each node. We regularly identified three communities, but in 17% of the time windows, we identified four communities. In 2% of the time windows, we identified only two communities with communities 2 and 3 merging. Even though node allegiance might change throughout the time course, the core of each community remained stable. The community centered at the right MTL and frontal regions in particular, shows very high stability.

**Supplementary Figure 5:**
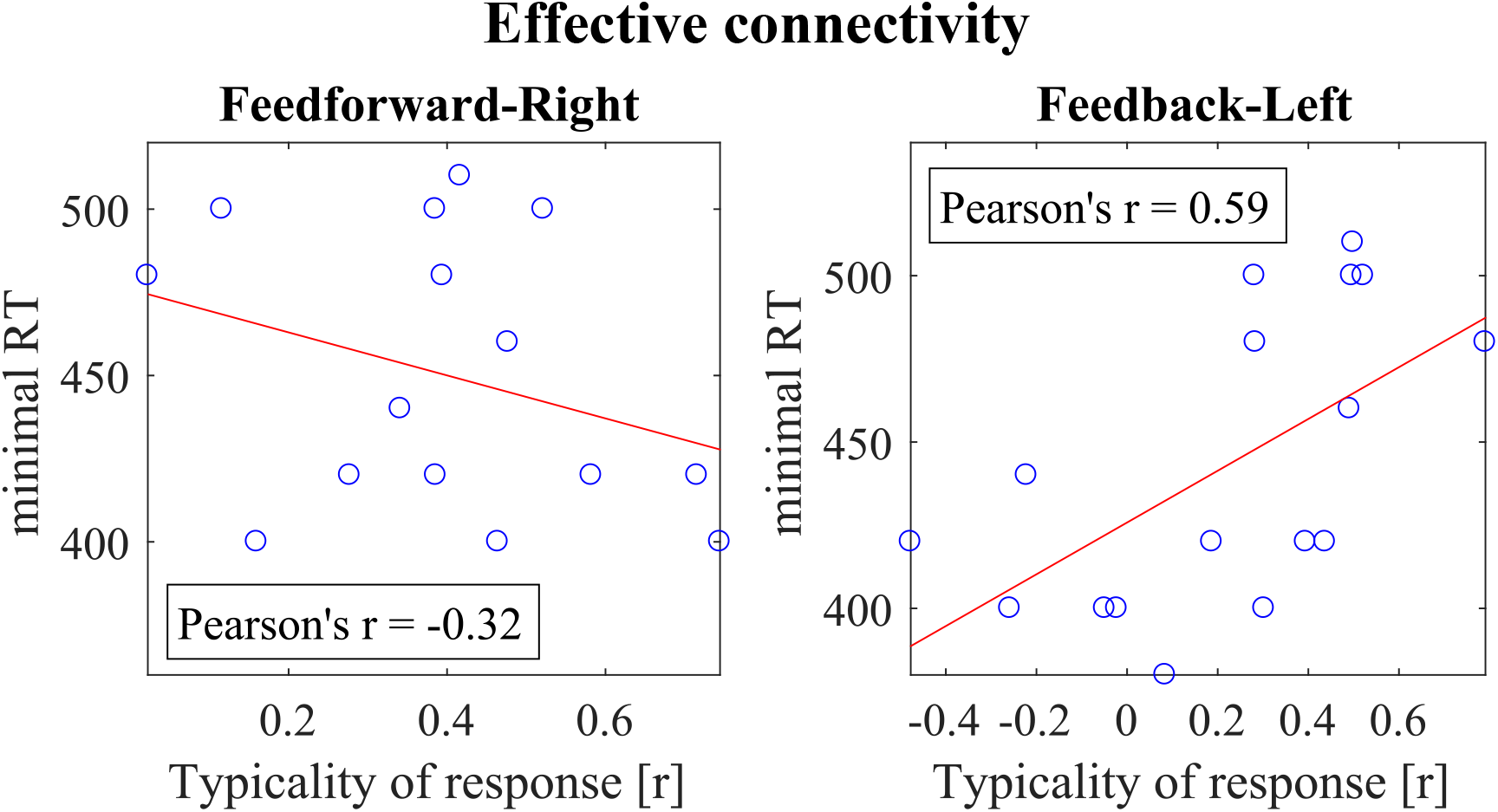
Each subject is described by the typicality of feedforward causality dynamics in the right hemisphere and feedback causality dynamics in the left hemisphere. We correlated the typicality with two measures of performance, namely the minimal reaction time and d-prime. Descriptively we observed that the feedforward connectivity was associated with fast responses and conversely for the feedback connectivity.

